# Optimal strategy for public good production is set by balancing intertemporal trade-off between population growth rate and carrying capacity

**DOI:** 10.1101/2021.07.22.453373

**Authors:** Manasi S. Gangan, Marcos M. Vasconcelos, Urbashi Mitra, Odilon Câmara, James Boedicker

## Abstract

Public goods are biomolecules that contribute to the community welfare. Their production can benefit populations in many ways, such as by providing access to previously unutilized resources. However, public good production has often been energetically costly, resulting in a reduction in the cellular growth rate. To reduce this cost, populations have evolved strategies to regulate biosynthesis of public good. Among these cell densities dependent regulation of public goods, as accomplished by quorum sensing, is a widely studied mechanism. Given that the fitness costs and benefits of public good production must be balanced, adoption of quorum sensing as a regulatory pathway by bacterial cells may have parallels with several economic principles that are used to study optimal investment decisions. Here, we explore the regulation of a public good, whose benefit is an increase in the carrying capacity, through experimental measurements of growth for engineered strains of *Escherichia coli* and analysis of those results using a modified logistic growth model. By varying the cell density at which the production of the public good was activated, we showed sharply-peaked optimum population fitness. Analysis further revealed that cell density associated with maximum public good benefits was determined by the trade-off between the cost of public good production, in terms of reduced growth rate, and benefits received from public good, in the form of increased carrying capacity. Moreover, our model showed that cells with *luxRI* quorum sensing seem to upregulate public good expression when the benefits from the production was immediate. These results demonstrate a case where a biological system apparently has evolved to optimize the timing of public good production to account for short-term costs and delays in reaping a future benefit.

**Author summary:** Bacteria often cooperate by sharing resources, sometimes referred as public goods. The activity of public good molecules benefits all of the neighboring cells. Here we examine the public good amylase, an enzyme to break down starch into smaller pieces that can be eaten by cells. Amylase production benefits the entire population because it enables growth to a larger population size, but production of this enzyme also has a cost. Enzyme production requires cells to expend energy and cellular resources. To balance the cost and benefit of the production of amylase, cells have likely evolved a strategy to control the timing of the production of public goods. We tested the dependence of population fitness on the timing of amylase production, finding an optimal cell density for amylase production. Finding the balance means that cells have evolved to anticipate future events, such as when simple food sources are depleted. Similar balancing of immediate costs and future benefits has been observed and studied in economics. Our results revealed that cells which use the regulatory strategy quorum sensing to control amylase production have minimized the delay in obtaining a benefit from this public good.

## Introduction

Often at higher cell densities, microorganisms favor cooperative interactions by producing metabolically expensive biomolecules known as public goods [1-3]. Important for growth and propagation, public good molecules are synthesized and secreted by cells into the extracellular environment and their activity is a benefit to both the producing cells and neighboring cells [4-6]. In the context of proliferating bacterial communities, public good supports continued growth and population stability, and delaying its synthesis can eventually decelerate population growth [7]. However, expression of public good imposes an immediate cost in terms of reduced intrinsic growth rate, in one example growth was reduced by as much as 83% [8], thereby affecting the net benefit of the public good. Production and utilization of public goods in microbial communities, thus, give rise to an economy, whose interest lies not only in maximizing the rewards delivered by public goods, but also reducing the energetic costs of public good production.

Economics recognizes this as a problem of optimization of trade-off between costs and benefits from production and consumption of goods. Theoretically, the populations that bear the consequences of public good production in real time are categorized in two groups. Populations that disregard future benefits from public good and prefer investing its resources to support growth in present are identified as impatient populations. On the other hand, populations that desire future gratification rather than present satisfaction are called patient populations [9]. Both populations seem to suffer losses in terms of stagnant growth in future or reduced intrinsic growth rate in present, respectively. Under such situations, economics explains that an ideal time for investments in public good production can be determined by analyzing the payoffs obtained from public good production and then evaluating the cost imposed due to delayed investments [10]. This raises the question how biological systems have evolved to account for the potential of a time-delayed benefit received from an energetically costly activity.

Generally, bacteria control public good expression through the process of quorum sensing [11, 12]. Quorum sensing involves the accumulation of an autoinducer signal, which in turn, enables cells to delay the production of public goods until reaching a high cell density [4]. Thus, populations of cells can use quorum sensing to coordinate the timing of public good production. Several studies have captured the importance of quorum sensing to tune public good production with cell density [8, 13-17]. However, its ability to negotiate current costs, such as an immediate reduction in growth rate, with future gains, such as an increased population, remains to be analyzed. Selective pressure should favor a regulatory mechanism to continuously probe population density over time to optimize the timing of the production of public goods. Here, we pose quorum sensing as a process that helps an individual bacterial cell to express public good at an ideal time and to extract a maximum net benefit.

To test this idea, we designed synthetic genetic circuit to quantify the time dependent variation in cost and benefit received from the expression of a public good. The public good, an α- amylase enzyme, increases the supply of nutrients via digestion of starch. The expression of the public good was timed by the exogenous addition of a high concentration of inducer, in this case the quorum sensing signal 3-oxo-C6-acylhomoserinelactone. Our experiments show that time is a key aspect in the formulation of a population decision of committing to public good expression. Under given growth conditions, expression of public good at an optimal time point balances the trade-off between population growth at present and increased carrying capacity in the future. We corroborate our experimental results with theoretical studies by proposing a modified logistic growth model for populations that synthesize energetically expensive biomolecules. In our model, the intrinsic growth rate and carrying capacity change over time according to an activation signal regulated by quorum sensing. Our experimental data shows that the switch between high and low growth rate and carrying capacity may not occur instantaneously. This potential latency in the change in the carrying capacity leads to an optimal density to activate α-amylase expression.

## Results

### Population density regulated expression of public goods optimizes the trade-off between population growth rate and carrying capacity

To experimentally test the fitness effects of public good production, we engineered a set of *Escherichia coli* strains to produce the public good α-amylase, an extracellular enzyme that digests polymeric starch into simpler sugars like glucose and maltose that can be absorbed and metabolized by the cells. The *amyE* gene encoding for α-amylase was copied from the genome of *Bacillus subtilis 168* and incorporated into a plasmid for heterologous expression in *E. coli* MG1655 (Fig S1a and S3). On the plasmid, public good was regulated by a quorum sensing gene circuit. Cells express both the *luxI* and *luxR* genes, which synthesize and detect the autoinducer molecule 3-oxo-C6-acylhomoserinelactone (AHL). The entire circuit as well as the public good gene were regulated by the quorum sensing-responsive *P*_*lux*_ promoter. Using this circuit, the production of α-amylase is delayed until the concentration of AHL exceeded a threshold concentration. Cells were grown in M9 media containing both starch and a low concentration glucose, such that cells utilizing only glucose would grow initially and sustained growth would require public good production. Once activated, cells expressed α-amylase (Fig S2a), which increased the amount of growth within the population.

To validate the basic cost and benefit associated with this production of the public good, we tested three growth strategies: OFF, ON and quorum sensing (QS), as shown in Figure 1a. In the ‘OFF’ strategy, *amyE* was deleted from the genetic circuit. ‘OFF’ populations cannot hydrolyze starch and only metabolize glucose. ‘ON’ cells contained the quorum sensing-regulated gene circuit and were stimulated through the addition of external AHL at the beginning of the experiment, such that the public good was produced continuously throughout the experiment. These cultures can consume free glucose and degrade starch as an additional food source. In the quorum sensing, ‘QS’, strategy, production of α-amylase was controlled via AHL autoinducer. As opposed to the ‘ON’ condition, external AHL was not added at the beginning of the experiment, but instead AHL was produced by the cells and accumulated in the culture medium over time. Quorum sensing was activated at cell density around 10^7^ CFU/ ml leading to public good production (Fig S4 a and b).

**Figure 1:**
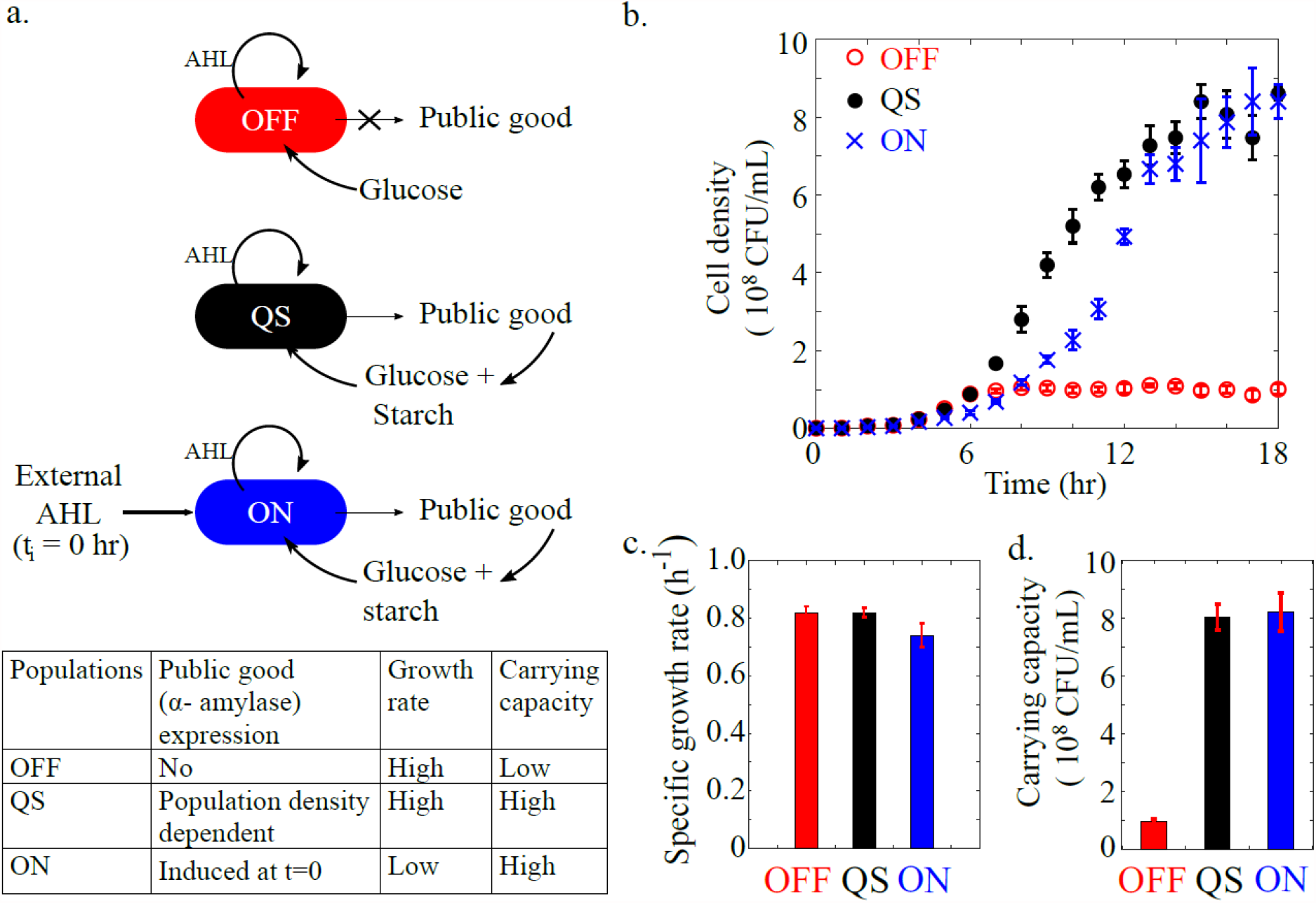
The cost and benefit of production of the public good α-amylase. (a) Depiction of three different strategies for production of a public good, α-amylase. OFF cells cannot produce the public good, QS (quorum sensing) produces public good at high cell density, and ON cells are externally induced to produce public good at the beginning of the experiment (t_i_ = 0). Cells are grown in media with glucose and starch as carbon sources, such that α-amylase increases the amount of carbon available for cell growth. (b) Growth of cultures in the conditions of OFF (red open circles), QS (black filled circles) and ON (blue cross) populations. (c) Specific growth rate between 0 and 6 hours and (d) Carrying capacities for the three regulatory strategies. For all plots n = 3 and error bars show standard deviation.

The growth behavior of these three strains, OFF, ON, and QS, was measured to determine the advantages and disadvantages of each strategy. All three strains were inoculated into M9 media supplemented with glucose and starch. Cell density was measured over time by spreading aliquots of culture onto LB agar plates and counting colony forming units. As shown in Figure 1b, growth dynamics differed between the three strains. The initial growth rates of ‘OFF’ and ‘QS’ were greater than the ‘ON’ population, indicating an energetic cost associated with public good production. There was also a change in the carrying capacity of these three strains. Growth was slowed down for the OFF strain at approximately 10^8^ cell/ml. The QS and ON strains continued to grow until approximately 8 x10^8^ cell/mL due to the ability of the public good to release additional nutrients from the media. These differences in the population growth patterns were quantified in terms of specific growth rate (Fig 1c) and carrying capacity (Fig 1d). The specific growth rate was calculated for these populations from 0 to 6 hours. The carrying capacity was calculated as the average cell density of the last three experimental time points. These metrics clarify the costs and benefits of public good production.

There is a clear cost to the public good, as demonstrated by reduced growth of the ON culture at early times. The benefit of amylase production under these culturing conditions is an increase in the carrying capacity, as shown in the ON and QS cultures. Our experiment demonstrated that for bacteria expending energy to gain an increased carrying capacity, the density-dependent regulation was a successful strategy to reduce the initial cost while maintaining the benefit of increased carrying capacity.

### The optimal delay to public good production maximizes population fitness

As shown in Figure 1, production of α-amylase has the benefit of increased carrying capacity and the cost of reduced growth rate. Regulating public good production via QS was more beneficial than both not producing the public good and always producing the public good. Next, we measured how the timing of public good production influenced population fitness.

To precisely regulate the timing of public good production, we created a new gene construct with a truncated *luxI* gene, thus cells could not produce the AHL inducer (Fig S1c and S5). Cells equipped with this circuit used glucose from the medium as carbon source until induction by externally added AHL. Induction led to expression of α-amylase and the extracellular digestion of starch into additional nutrients that could be used for growth (Fig 2a and S2c). In parallel cultures, production of the public good α-amylase was induced at different times (t_i_), and changes in cell density were monitored by plate counts.

**Figure 2:**
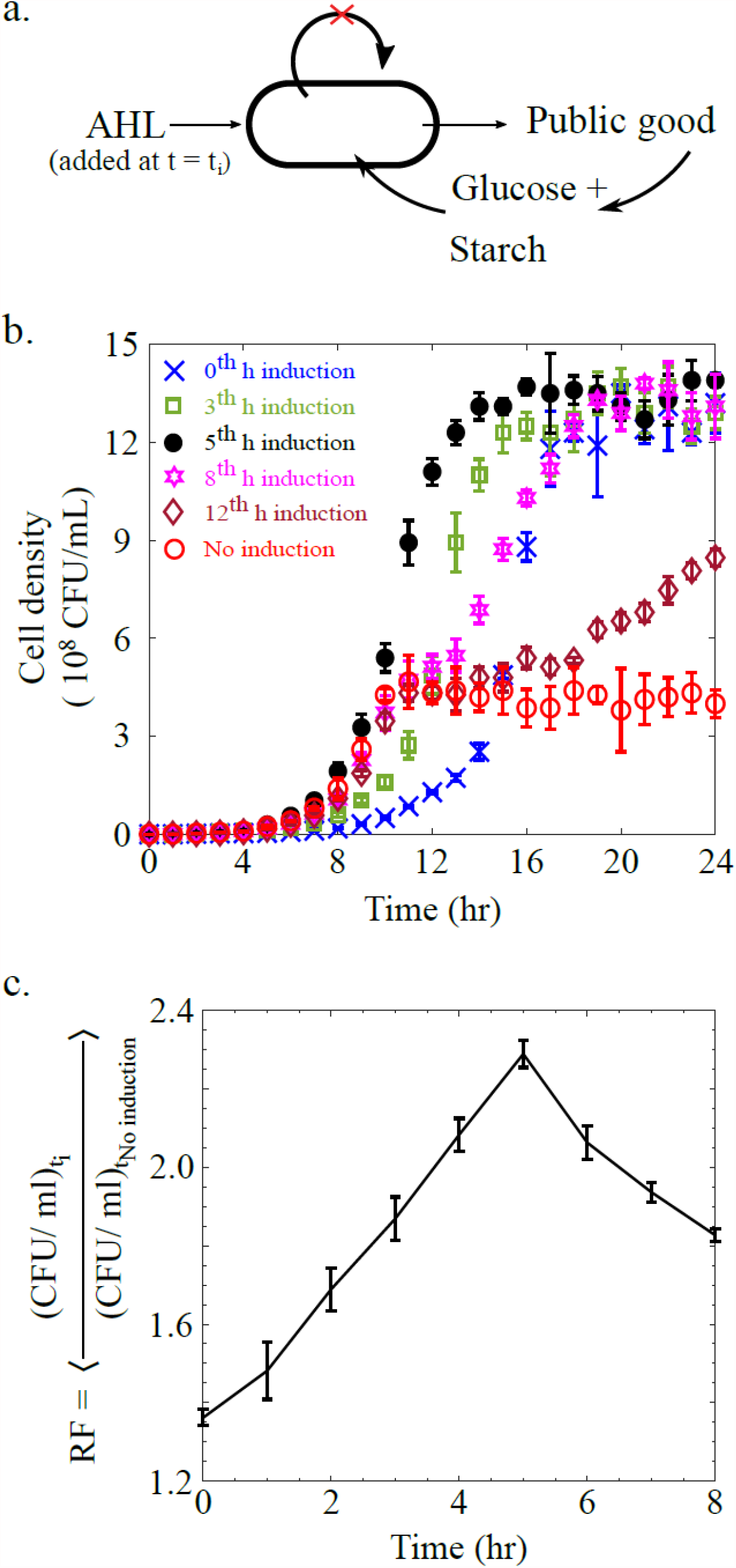
Finding the optimal time to initiate public good production. (a) Schematic shows public good production when induced by external AHL at t = t_i_. Cells in these experiments do not synthesize AHL. (b) Growth of individual populations activated with AHL added externally at 0, 3, 5, 8, and 12 hours. No induction shown as negative control. (c) Fitness of the induced cultures relative to the uninduced culture averaged over the entire experiment. The relative fitness is plotted as a function of the induction time. n = 3 for all measurements and error bars show standard deviation.

As shown in Figure 2b, cultures that were induced at early times had slower initial growth, yet all induced cultures eventually reached the elevated carrying capacity. As the induction time was delayed, cultures maintained the higher specific growth rate for longer times. See Fig S6a for data from additional induction times. To find the optimal time to initiate production of α-amylase, the relative fitness of each culture was calculated (Fig 2c and Fig S6b). Relative fitness (RF) is the ratio of cell density of a given culture to the uninduced negative control averaged over the entire experiment, from 0 to 24 hours. The maximum relative fitness was for induction at 5 hours. Our results substantiate that an optimum time delay preceding the synthesis of a public good, maximized cell fitness. Next, a mathematical model explored how adjustment of the threshold parameter, the density at which activation occurred, influences cell growth and leads to an optimal time to initiate public good production.

### Developing a model for public good production

Our mathematical model is a modified version of the traditional logistic growth model:

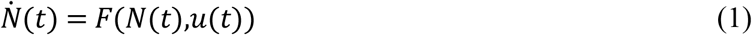

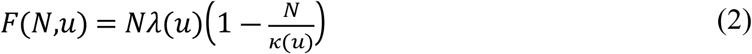

where N(t) denotes the cell density at time t. The logistic growth model is characterized by two parameters: the intrinsic growth rate (*λ*), and the carrying capacity (*κ*). As depicted in Fig 3a, the control signal that modifies the growth rate and carrying capacity of the colony is denoted by *u*. The control signal is a function of the concentration of the AHL autoinducer used to regulate expression of quorum sensing responsive genes, here the public-good α-amylase. For the measurements with the ‘QS’ strain in Fig 1 this signal accumulates over time as a result of AHL synthesis by the cells. For the ‘ON’ strain in Fig 1 and the externally induced strains used in Fig 2a, a high concentration of signal was added at time *t*_*i*_.

**Figure 3:**
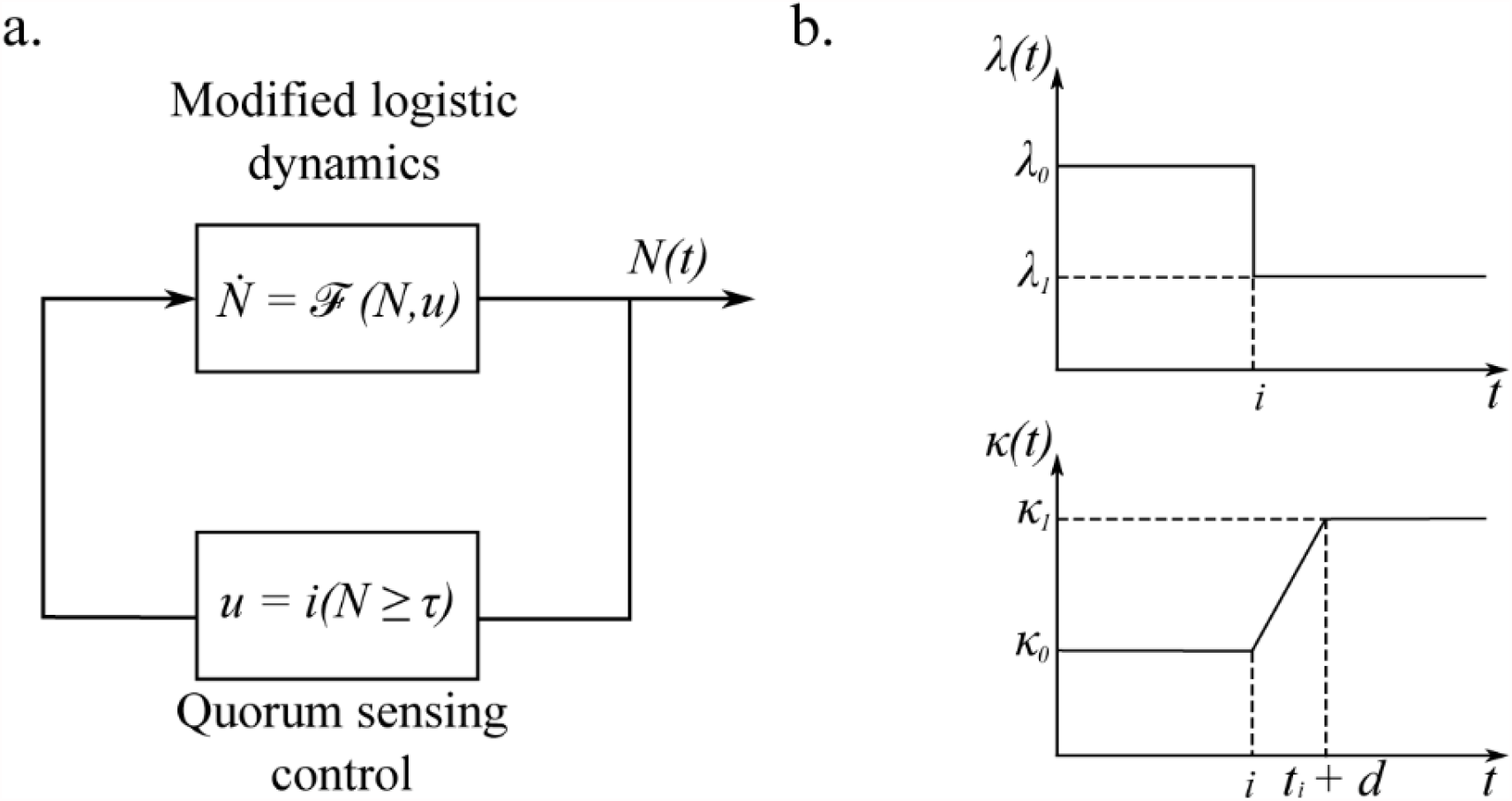
Developing a modified logistic growth equation to incorporate the cost and benefit of public good production. (a) Block diagram of our system model for control of bacterial growth via quorum sensing. The state of the system is the number of cells, *N*. The control signal, *u*, corresponds to the decision of activating or not the production of a costly enzyme. The policy used to compute the control implements quorum sensing, where activation occurs once a certain target population is met. (b) Changes in the growth rate and carrying capacity over time. The decision to produce the public good is reached at t = i.

In the model, activation of quorum sensing, beyond time *t*_*i*_ or when the AHL concentration exceeds a threshold concentration for the ‘QS’ strain, results in a switch in the value of both the carrying capacity and the specific growth rate. The growth rate is decreased due to the burden of public good production and the carrying capacity is increased as additional nutrients are now available for growth. We assume that the switch in the growth rate constant is instantaneous, as depicted in Fig 3b. The increase in the carrying capacity is not always instantaneous, as suggested by the results shown in Fig 3b when external signal was added at t = 12h. Before 12h, the culture had already reached the carrying capacity, and activation resulted in a slow drift of the population size from the lower carrying capacity towards the higher carrying capacity. The scheme for the gradual change in the carrying capacity is depicted in Fig 3b and incorporated into the model using:

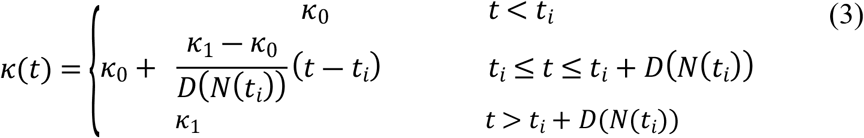

*D* accounts for the time it takes to change from the smaller carrying capacity to the larger carrying capacity. This delay may have many contributions, including the time needed for enzyme production and sufficient starch degradation to impact cell growth.

If there is no delay (*D*(*N*(*t*_*i*_)) = 0), then our model for a quorum sensing population simplifies to:

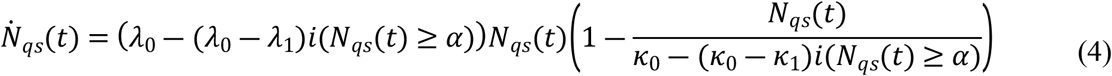

Where the function *i*(*S*) denotes the indicator function of the argument S, i.e., *i*(*S*) = 1 if *S* is true, and *i*(*S*) = 0 if *S* is false.

### Fitting experimental data to calculate the delay in the benefit from public good production

Next, we implemented the model to calculate the values of the parameters in Equations 1-3. Figure 2 reports a set of 3 independent time series data for the population activated at times *t*_*i*_ ranging from 0 to 12h, including a culture that was not activated. Each series contains 25 samples, taken in 1-hour long intervals. For simplicity, we have used a discrete-time approximation of the model in Equations 1-3. To calibrate the model, the first step is to estimate the carrying capacity and the intrinsic growth rates for the ON and OFF cultures, t_i_ = 0 and no induction, respectively. This was done using a weighted non-linear least squares regression, in which we fixed the initial condition of our model *N*(0) as the average of the first sample in each of the three data series corresponding to the ON or OFF cultures. This step is necessary because the data varies by several orders of magnitude from the first sample to the last. Therefore, with a free initial condition as an optimization parameter, least-squares will ascribe a much larger weight to the steady-state regime, which leads to very good estimates of the carrying capacity, and poor estimates of the intrinsic-growth rate. Similarly, a least-squares optimization in log-scale, creates the opposite problem resulting in an excellent estimate for the growth rate, but a poor estimate for the carrying capacity. Albeit atypical, the strategy of fixing the initial condition and performing regular least-squares fitting leads to good estimates for both quantities. Also notice that the growth rate and capacity were obtained for the discrete-time model, which was compared with the sample average of the data using least-squares as follows:

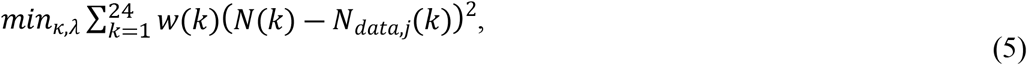

where the weights *w*(*k*) are the inverse of the standard deviation of the measurements *N*_*data,j*_(*k*), *j* = 1,2,3.

Best-fit values for *κ* and *λ* for the ON and OFF cultures are shown in Fig 4. The best-fit value of the specific growth rate for the OFF culture is much higher than the previously calculated value reported in Fig 1c, as Fig 1c shows the results of an exponential fit to the first 6 timepoints and Fig 4 shows the results from a fit to the logistic growth equation.

**Figure 4:**
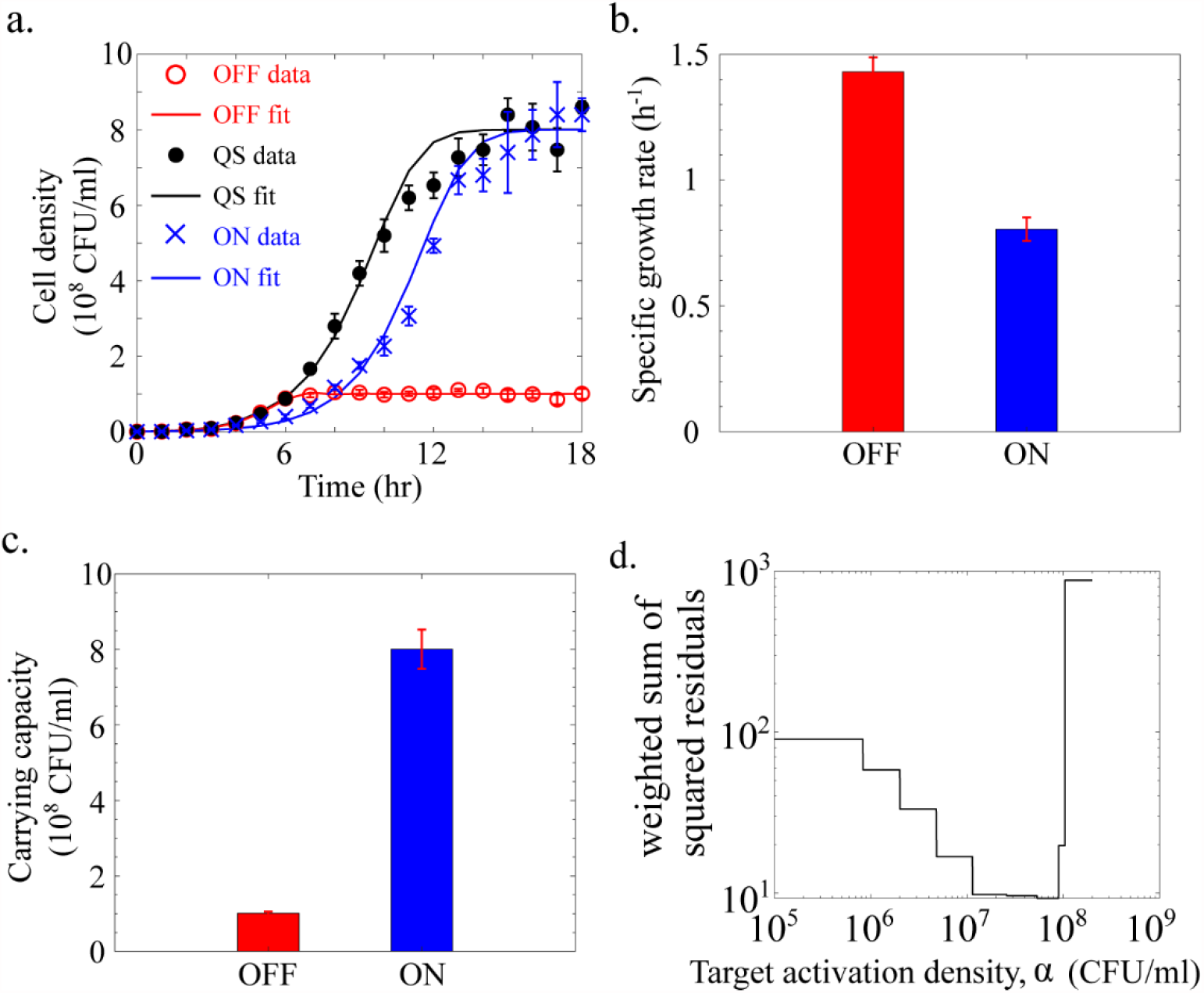
Fits to the modified logistic growth equation. (a-c) Using the experimental data from the ON and OFF strategies reported in Fig 1B, best-fit values for the *λ*_0_, *λ*_1_, *κ*_0_ and *κ*_1_were calculated. The values for these parameters were used to predict the growth dynamics of the strain following the quorum sensing strategy described in Eq. (4). To find the target activation density *α*^∗^, we solve another nonlinear weighted least squares regression problem. For the quorum sensing strain optimal activation threshold *α*^∗^= 5.3 10^7^ CFU/mL. (d) Weighted sum of the squared residuals as a function of *α*.

The second step in the data analysis was to estimate the delay in reaching the gain in carrying capacity for each of the induction times *t*_*i*_. The procedure we used was slightly different than the one described above. For each activation time *t*_*i*_, we perform a non-linear least-squares optimization over the delay parameter *D*, from Eq 3, and the initial condition *N*(0), as follows:

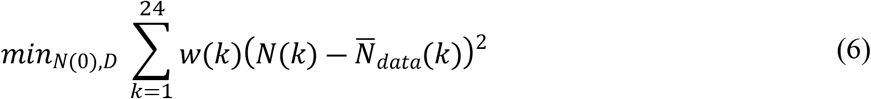

where *N*(*t*) is computed according to a discrete-time approximation of Equations (1-3), and w(k) is a weight which is inversely proportional to the standard deviation of the time-series data.

The reason why the joint optimization of *N*(0) and *D* works well in this case is that the growth rates and carrying capacities (*λ*_0_, *λ*_1_, *κ*_0_, *κ*_1_) are now fixed, and the issue of the transient and stationary regimes dominating the objective function does not exist in this case. Using the best-fit values of the delay shown in Fig 5a, Figure 5c compares the experimental data and to the model predictions.

**Figure 5:**
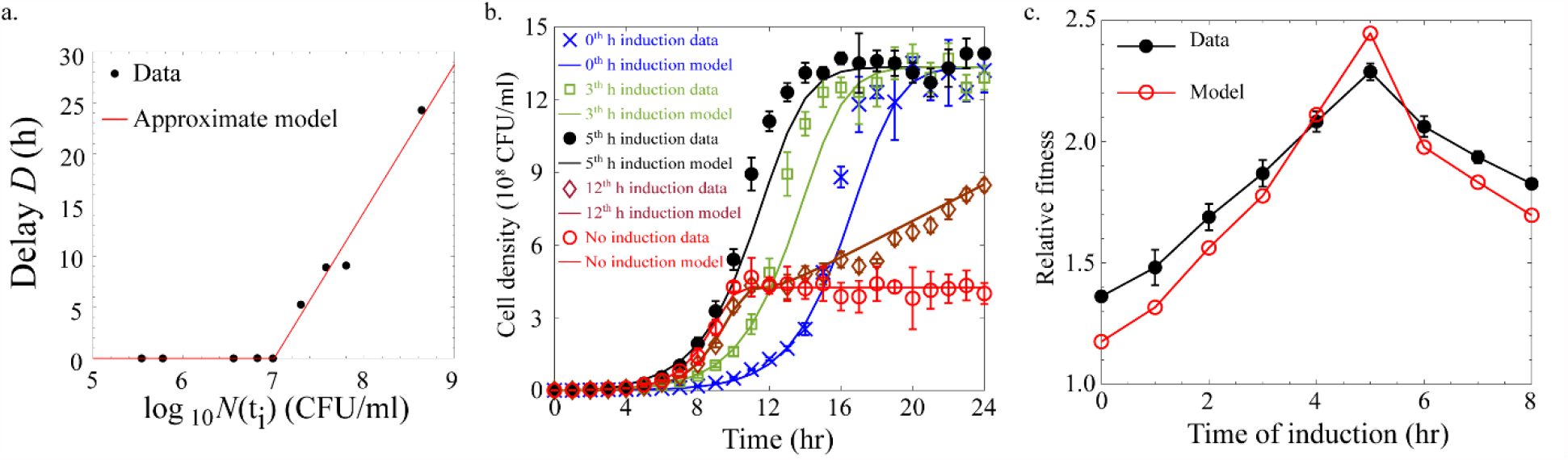
Fits for the delay in the benefit from public good production. (a) From the experimental data, the best-fit value of the delay time was calculated for each induction time t_i_. The red line is the piece-wise linear function describing the relationship between delay in carrying capacity as a function of the population at the activation time (b) The solid lines show the model prediction made after fitting for the delay parameter *D*. Symbols show experimental measurements. (c) Empirical and theoretical fitness function computed from data and our calibrated mathematical model, respectively, which shows that our simple model (with a minimal number of parameters) successfully captures the fundamental trade-off that occurs in bacterial growth controlled via QS.

As shown in Fig 5a, the highest population density at which there is no delay in receiving the public benefit from public good production is at approximately 1 × 10^7^ CFU/mL. There is a delay in the change in the carrying capacity if α-amylase production is initiated at a cell densities above this range. As shown in Fig 5d, when using the best-fit parameters for the growth rates and carrying capacities and the delays in public good benefit for each activation density, we reproduce the relative fitness trend calculated from the experimental data. There is a density at which the initiation of public good production optimizes fitness, and this optimal density corresponds to a density for which there is no delay in the public good benefit. The penalty, or delay of the public good benefit, increased for cultures that initiated public good production further outside of this optimal range. These findings demonstrate that for α-amylase production as a public good, there is an optimal range of cell densities to initiate public good production. Nonlinear least-squares regression of the data for the quorum sensing strain in Fig 1 (‘QS’) revealed activation of quorum sensing in the range of 5∼9 × 10^7^ CFU/mL, which is very close to the optimal strategy for regulation of public good production.

## Discussion

In economics, decisions with consequences that play out in future, are described by the concept of intertemporal choices [18]. When making a decision, individuals tend to assign an additional cost to rewards received at a later time [9, 10, 19]. This creates a specific point in time at which benefits of the decision exceed its cost, thereby maximizing the trade-off between the two. Personal goals of a decision maker play crucial role in determining the ideal time for action [20].

The model considered herein captures the fundamental trade-offs observed in the experimental data. Importantly, it also accounts for the latency observed during activation past the typical values of activation observed in nature. One major distinction of our model with previous work in this area is that a reduction in growth rate leads to a gain in capacity. Previous works have not explored a switch in carrying capacity, see for example the work of Pai et al [8].

For a bacterial cell, the decision to expend energy for public good production also has an intertemporal dimension. Cultures that activate too early bear the cost of public good production before these resources are needed, whereas waiting too long to produce public goods leads to diauxic growth, with a lag between glucose and starch utilization. Early production of public good at lower cell densities, shunts cellular energy into biochemical pathways rather than growth and reproduction. As a result, public good production compromises the instantaneous growth rate of the population. However, the strategy to stall public good production to support the current population growth rate, leaves populations unprepared for future growth conditions and eventually has a negative effect on population fitness [7, 8, 13, 21]. Postponing the expression reduces glucose content of the environment and reduces the growth until glucose is again made available. Here, we have studied quorum sensing as a process to set the intrinsic population growth rate while deciding an appropriate time for an activation of public good production, thereby optimizing the trade-off between population growth rate and fitness continuously in time.

Earlier studies that discuss the trade-off between population growth rate and carrying capacity, emphasize on the tendency of the populations to use catabolic pathways associated either with fast but lower ATP output, inefficient catabolism, or with slower but higher ATP yield, efficient catabolism [22-24]. Since, population growth is dependent on ATP production, use of either of the metabolic pathway impacts the trade-off. Inefficient ATP production has been shown to increase population growth rate but reduces the biomass yield, while efficient ATP production is known to hamper the growth rate but eventually increases the carrying capacity [23]. Thus, these studies establish the role of ATP output as deciding factor in trade-off between population growth rate and hint at the evolution of biochemical pathways to guide this trade-off. However, the existence of a biochemical pathway to optimize the trade-off between instantaneous population growth rate and carrying capacity has never been explored.

Here, modeling of the transition to the utilization of starch as a nutrient has quantified the delay in a population reaping the benefit of the public good α-amylase. The model was adapted from previous work on quorum sensing as an optimal control system. Previous work has shown that under the assumption of perfect observation of the state of the system, namely, the population at a given time, a threshold policy on the population maximizes a discounted objective function. Later, this discrete time model [25] was extended to continuous time [26]. More importantly, while assuming a more realistic model with partial observations of the state of the colony via realistic AHL signal dynamics, the discounted objective function was numerically shown to be unimodal in the target population threshold. Similarly, precise timing of public good production was shown here to optimize fitness. If a population activates α-amylase expression at too low of a cell density, there is a delay in the time between α-amylase production and the time at which the population requires starch degradation to maintain growth. There is also a delay in the benefit from the public good if production of α-amylase occurs after growth has begun to slow. This delay represents a lag in the ability of cells to metabolize starch degradation products, and the lag increases as public good production is further delayed. These two delays in realizing the benefit from α-amylase production creates an optimal cell density of public good production.

Over the years several studies have considered the importance of quorum sensing as a control mechanism for public good production [7, 8, 13-15, 27-29]. Work done by You and colleagues has used a similar framework to establish the optimal regulation of quorum sensing. This study follows the growth of synthetic strain of *E. coli* that expresses public good to survive under antibiotic stress. Though this work does not discuss the changes in the carrying capacity with public good production, it shows that the optimal population growth rate is possible only when public good production is switched on through quorum sensing [8]. In our system, the cost of production of public good is reflected in the growth rate and the public benefit is the net gain in carrying capacity. Growth models where the individuals control the colony’s carrying capacity were previously studied in the social sciences in the work of Meyer and Ausubel [30], where the adoption of a new strategy may create newly available resources that allow for higher population yield. Analogously to the bacterial case at hand, the adoption of a new strategy is a gradual process and does not yield immediate gains in carrying capacity. In other words, there is an inherent delay, especially if the population is already large. The work of de Vos et al [31]. considered the how interactions between different strains in a polymicrobial infection affect the growth rate and the carrying capacity. However, this change in behavior was passive rather than active, as in our model.

In conclusion, here a bacterial population was shown to optimize the timing of public good production, such that there is no delay in receiving the benefit from the public good amylase. This optimal timing accounts weighs the short-term changes in the intrinsic growth rate with the long-term goal of maximizing the carrying capacity. Comparison of the regulatory strategies for public good production in different bacterial species may reveal how individual species have navigated such trade-offs in short-term and long-term costs and benefits. The cost and benefit of specific public goods should vary and will depend on environmental conditions, so perhaps such regulatory strategies will have evolved to be beneficial for typical environmental contexts. Certainly, evolutionary forces have the ability to adjust parameters such as the threshold signal concentration, or likewise the cell density of achieving a quorum. Incorporating the potential for reduction for future benefits due to time delays into such analysis may reveal new aspects of how populations of cells anticipate future conditions and evolve optimal regulatory strategies.

## Materials and Methods

### Strains and culturing conditions

*E. coli* MG1655 cells [32] were used as a parent strain for the propagation and expression of variations of the genetic circuit. Cultures were grown at 37 ° C at 200 rpm in M9 media supplemented with a carbon source and 50 µg/ml Kanamycin for plasmid maintenance. Primary cultures were grown overnight in M9 (Difco™) media supplemented with 4% glucose (aMResco®).

### Plasmid construction

Genetic circuits for this study were designed by modifying pTD103*luxI*_*sfGFP* plasmid [33] to test three strategies of α-amylase production that included ON (constitutive expression), QS (density-dependent expression) and OFF (no expression). Gibson assembly was used for plasmid construction (NEB). *amyE* gene encoding for α-amylase was amplified from genomic DNA of *B. subtilis 168* (ATCC®23857™) and cloned into pTD103*luxI*_*sfGFP* under the regulation of the *P*_*lux*_ promoter, creating plasmid pPG_*amyE*^*+*^ used for measurements of the ON/QS strategy. The plasmid used for the OFF strategy, pPG_*amyE*^*-*^ was created by deleting the *amyE* ORF from plasmid pPG_*amyE*^*+*^. See Fig S1a and S1b.

For experiments to test the timing of public good production on growth dynamics, *luxI* gene was partially deleted from plasmid pPG_*amyE*^*+*^ to create plasmid pPG_*amyE*^*+*^_*ΔluxI*, to deactivate acetyl transferase activity of LuxI protein [34]. Strains harboring this plasmid can respond to but do not synthesize the QS signal, therefore activation of public good production was induced by adding an external stimulus of signal. See Figs S1c and S5.

### Growth Conditions

#### Optimization of trade-off between population growth rate and carrying capacity

To test the growth of cells using the ON, OFF, and QS strategies, primary cultures were diluted 1000X in 5 ml fresh media. For ON cells, public good production was induced by adding 3 µg/ml of 3-oxo-C6-acylhomoserinelactone (AHL, Adipogen) as in [35]. Upon reaching OD_600nm_ of 0.3, cells were washed three times 1X PBS (VWR, life science) and resuspended in 1 ml 1X PBS. Cells at 100X dilution were then inoculated into 5 ml fresh M9 media supplemented with 0.0125% glucose and 0.5% starch (Sigma-Aldrich). For the ON cells, 3 µg/ml of AHL was used to induce public good production. The cell density within triplicate cultures was monitored every hour for 18 hrs by counting colony forming units on LB agar with 50 µg/ml kanamycin.

#### Time dependent activation of public good

Primary and secondary cultures of cells harboring plasmid pPG_*amyE*^*+*^_*ΔluxI* were grown using the procedure for the QS strategy. Cells from the secondary culture were used to inoculate M9 media supplemented with 0.0125% glucose and 0.5% starch at 100X dilution. Using this timepoint as t=0, 3 µg/ml of 3-oxo-C6-acylhomoserinelactone was added at the induction time. For cells labeled “no induction”, no external signal was added. The cell density within triplicate cultures was monitored every hour for 24 hrs by counting colony forming units on LB agar with 50 µg/ml kanamycin. For these experiments, cells induced at t=0 were prepared following the procedure for the ON strategy described above.

### Analysis

Specific growth rate of bacterial populations was measured by fitting a straight line to the natural log of cell density over time for data between t=0 and t=6 h.

Carrying capacity was calculated by taking average of CFU/ml of the last three time points.

To calculate relative fitness of an induced cultures, the mean cell density of the induced culture over 24 hrs was divided by the mean cell density of the uninduced cultures, using:

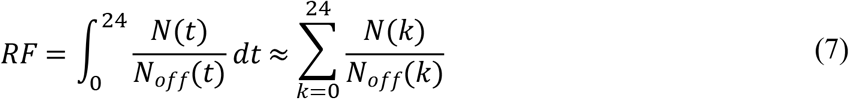

This definition is appropriate because despite the fact that the transient and stationary regimes are orders of magnitude apart, they are cancelled in the relative ratio inside the integral. Therefore, the integrand has approximately the same order of magnitude across time.

## Acknowledgements

JB, UM, MV, and MG acknowledge support from Army Research Office MURI Award W911NF1910269.

## Supplementary Information

### Supplementary methods

Confirmation of α-amylase production and activity

Quantification of 3-oxo-C6-acyl homoserine lactone production by pPG_*amyE*^*+*^ cells Quantification of LuxI activity.

### Supplementary figures and tables

Table S1: Plasmids used in this study.

Figure S1: Plasmid maps.

Figure S2: Schematic representations of public good regulatory strategies.

Figure S3: Expression of α-amylase.

Figure S4: Signal production and amylase activity over time.

Figure S5: *luxI* mutant does not induce expression of quorum sensing regulated genes.

Figure S6: Extended data set for public good production induced between 0 and 12 hours.

